# Pharmacological Stratification of Public Bioactivity Databases: A Reusable, OECD-Anchored Curation and Benchmarking Framework Demonstrated for Opioid Receptors

**DOI:** 10.64898/2026.06.18.732083

**Authors:** Manal A. Nael, Laxman M. Alakonda, Abhik Ghosh, Sara Jane Ward, Lee-Yuan Liu Chen, Anjali M. Rajadhyaksha, Magid Abou-Gharbia, Khaled M. Elokely

## Abstract

Public bioactivity databases are heterogeneous not only in measurement type, where binding affinities and functional potencies are reported on different scales, but in pharmacology: the same compound and target can carry agonist, antagonist, or inhibitor records measured through binding displacement, cAMP, β-arrestin, or [^35^S]GTPγS readouts that quantify different biological events. Pooling these records produces models whose output is detached from any coherent pharmacological claim. Prior work has standardized bioactivity at scale and quantified the noise from mixing measurement types, but pharmacological mechanism and assay-readout class have not been treated as a primary axis of large-scale curation. This study presents an auditable, OECD-anchored framework that stratifies public records by action type and assay readout before modeling, converting heterogeneous data into externally validated, interpretable QSAR tasks that compose with existing standardization resources rather than replacing them. The framework is demonstrated on the four opioid receptors (MOR, DOR, KOR, and nociceptin/orphanin FQ, NOP). Four public sources were reconciled into 72,148 merged records and 50,977 curated measurements spanning 19,585 compounds, each carrying auditable attributes for source agreement, endpoint meaning, pharmacology class, assay readout, and trust tier. Receptor-level binding tasks formed a compact benchmark with strong locked external performance, including KOR pKᵢ (R^2^ = 0.79, n = 798) and DOR pKᵢ (R^2^ = 0.77, n = 736). Pharmacology- and readout-resolved functional endpoints yielded externally validated strata that pooled labels would obscure, including a MOR antagonist functional-inhibition endpoint (R^2^ = 0.86, n = 110) and agonist potency endpoints for DOR, KOR, and MOR (R^2^ up to 0.81). Comparison against a fully pooled baseline shows that pooled models either match stratified models on coherent endpoints or reach a deceptively high R^2^ on functional-IC_50_ endpoints by training predominantly on binding-displacement records, so the pooled number predicts affinity rather than functional activity. SHAP attribution indicates that binding and functional potency encode partially distinct structure-activity signals. The dataset contract, not model performance alone, defines the validity and scope of a QSAR claim, and stratification is a precondition for a functional model to support a defensible claim. Curation logic, derived tables, frozen data, and reproducibility artifacts are released.

## Introduction

Public bioactivity databases have made it possible to build quantitative structure-activity relationship (QSAR) benchmarks at a scale that would be impractical for any single laboratory.^(1–5)^ These resources aggregate decades of medicinal chemistry and pharmacology across thousands of targets, and they increasingly serve as the default evidence base for machine-learning models of molecular activity.^(6)^ The promise of this scale is that larger, more diverse training data should yield models that generalize. The difficulty is that the value of any such model is bounded by how the underlying records are curated before training: a prediction is only as well defined as the endpoint, evidence tier, and validation context that produced it.^(7–12)^

One dimension of this problem is now well recognized. Public records are not generated under a single assay protocol, unit convention, or activity definition, and combining measurements of different types introduces measurable noise.^(13, 14)^ Systematic analyses have shown that pooling IC_50_ or K*_i_* values from different sources introduces errors large enough to limit model quality, with a substantial fraction of paired measurements differing by more than an order of magnitude.^(15–19)^ In response, large-scale standardization efforts have harmonized structures, units, and activity types across millions of datapoints to produce machine-learning-ready bioactivity collections.^(20, 21)^ These efforts address the measurement-type and chemical-representation axes of heterogeneity and provide an essential foundation for data-driven modeling.

A second axis of heterogeneity is not addressed by measurement-type harmonization or by standardization at scale, and it is the focus of this study. The same target can be characterized by records that differ in *pharmacological mechanism* and in *assay readout*. An agonist EC_50_ quantifies the concentration required for half-maximal receptor activation; an antagonist or inhibitor potency quantifies blockade of a reference response, suppression of signaling, or competition in a binding assay. A cAMP assay, a [^35^S]GTPγS assay, a β-arrestin recruitment assay, and a radioligand displacement assay report different biological events for the same compound, and the differences are pharmacologically meaningful rather than random error.^(22–26)^ Pooling agonist with antagonist data, or binding with functional readouts, is therefore not a units problem to be normalized away; it is a biology problem that forces a model to learn incompatible structure-activity relationships. To our knowledge, no large-scale curation of public bioactivity databases has treated pharmacological mechanism and readout class as a primary, systematic stratification axis applied before modeling. This study introduces that axis as a curation methodology, demonstrates that it changes which QSAR claims are defensible, and shows that it composes with, rather than competes with, existing standardization resources.

These considerations converge on a single principle: defining the dataset defines the QSAR claim.^(27)^ A model trained on records that mix binding and function, agonism and antagonism, or concordant and discordant source evidence may still emit numerical predictions, but the scientific claim those predictions support becomes ambiguous. Conversely, a smaller but pharmacologically defined endpoint can support a stronger claim when its provenance, chemical identity, activity scale, assay readout, pharmacology class, and external validation are made explicit. This principle was advanced conceptually in a prior Viewpoint;^(27)^ the present study is its empirical, large-scale realization, operationalizing the principle as a reusable, auditable curation framework whose outputs are inspectable at the level of individual records.

The opioid receptor family is an ideal stress test for this framework. MOR, DOR, KOR, and NOP form one of the most consequential G-protein-coupled receptor (GPCR) systems in drug discovery, central to analgesia, addiction, respiratory depression, tolerance, and the pursuit of pathway-selective ligands.^(28)^ They are supported by dense, multi-source data spanning binding and functional assays, and the agonist-antagonist and biased-signaling distinctions among opioid ligands have direct biological consequence. If pharmacological stratification matters anywhere, it matters here. Applying the framework to these four receptors, this study contributes: (i) a reusable pharmacological- and readout-stratification curation layer for public bioactivity data; (ii) empirical evidence that this stratification changes which QSAR claims are externally defensible; (iii) externally validated binding and functional benchmarks across the receptor family; (iv) an interpretability layer indicating partially distinct structure-activity signals for binding and function; and (v) released curation logic, derived tables, and frozen data for independent reproduction and reuse.

## Results

### A dataset contract defines the QSAR claim

The central proposition of this study is general: a QSAR claim is not defined at the model-training step but earlier, by the dataset contract that determines what each compound represents, which receptor and endpoint are modeled, whether source records agree, how assay context is interpreted, which records are trusted, and what form of external generalization is evaluated. Figure 1 illustrates this architecture for the opioid receptor demonstration. Four public sources were integrated into a single bioactivity resource, yielding 115,232 raw rows before reconciliation and curation. These were transformed into 72,148 merged records, 50,977 curated measurements, and 19,585 final compounds. The resulting endpoint matrices separate receptor-level binding tasks from functional tasks defined by pharmacology class and assay readout. A per-source contribution breakdown is provided in Tables S2 and S3.

**Figure 1.**
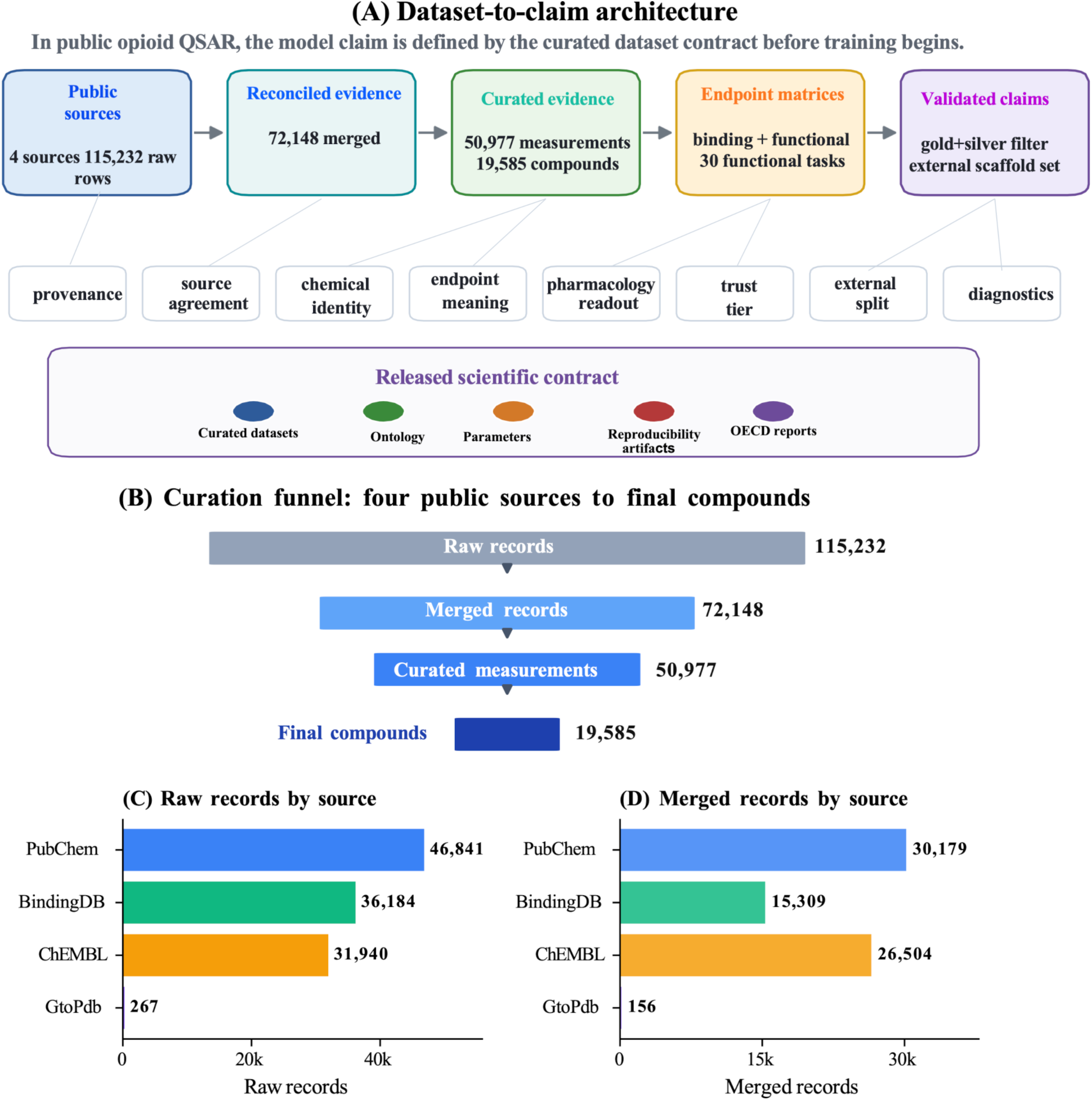
Dataset-to-claim architecture and curation funnel for bioactivity benchmarking, demonstrated on opioid receptors. (A) In public opioid QSAR, the claim a model can support is defined by the curated dataset contract before training begins. Public records pass through five stages, from public sources to reconciled evidence, curated evidence, endpoint matrices, and validated claims, gated at each step by explicit credibility checks for provenance, source agreement, chemical identity, endpoint meaning, pharmacology and readout annotation, trust tier, external splitting, and validation diagnostics. The released scientific contract comprises curated datasets, ontology, parameters, reproducibility artifacts, and OECD reports, so that the model is downstream of the curation logic and the dataset definition governs what the model is allowed to claim. (B) Curation funnel: four public sources contributed 115,232 raw records, reconciled into 72,148 merged records and 50,977 curated measurements across 19,585 compounds. (C) Raw record contribution by source and (D) merged record contribution by source, across ChEMBL, BindingDB, PubChem, and the IUPHAR/BPS Guide to Pharmacology (GtoPdb).

The first layer of the contract defines the evidence universe. Records were collected for MOR, DOR, KOR, and NOP from ChEMBL, BindingDB, PubChem, and the IUPHAR/BPS Guide to Pharmacology (GtoPdb).^(1, 2, 29, 30)^ Multi-source retrieval increases coverage but introduces overlap, annotation differences, and potential disagreement,^(31, 32)^ so records were not concatenated. They were reconciled into source-supported evidence groups with agreement and disagreement preserved as explicit metadata, because a repeated record is not automatically independent evidence; it becomes stronger evidence only when source annotations and values are concordant.

The second layer defines chemical identity. Because scaffold splitting, duplicate detection, and descriptor generation all depend on stable identity, deterministic chemical standardization was applied before endpoint aggregation. The third layer defines endpoint meaning: binding affinity, binding-displacement potency, agonist and antagonist potency, inhibitor-like responses, efficacy, and pathway-specific functional readouts are not one activity label, so endpoint meaning, pharmacology class, and assay readout are placed as separate credibility gates before matrix construction. The fourth layer defines evidence quality through trust tiers, retaining gold and silver records for the primary benchmark while keeping exploratory or unresolved records visible for audit. The fifth layer defines the validation claim through a scaffold-locked external set, so that reported performance tests chemical generalization rather than analog interpolation. As shown in Figure 1, the released contract includes the curated datasets, the pharmacology and assay-readout ontology, and the reproducibility artifacts; the model is downstream of the curation logic, and the dataset definition governs what the model is allowed to claim.

### Curation converts heterogeneous public records into auditable evidence

Curation does more than reduce row counts; it changes the evidentiary structure of the data (Figure 2). The compression path begins with 115,232 raw records and proceeds to 72,148 merged records, 50,977 curated measurements, and 19,585 final compounds. Alongside the retained evidence, the workflow records 14,116 explicit exclusions, 18,869 development-set review-queue records (3,456 in the external set), and 57,313 decision-lineage entries. These sidecar outputs document how the curated benchmark was produced, so that retained, excluded, and review-needed records remain traceable.

**Figure 2.**
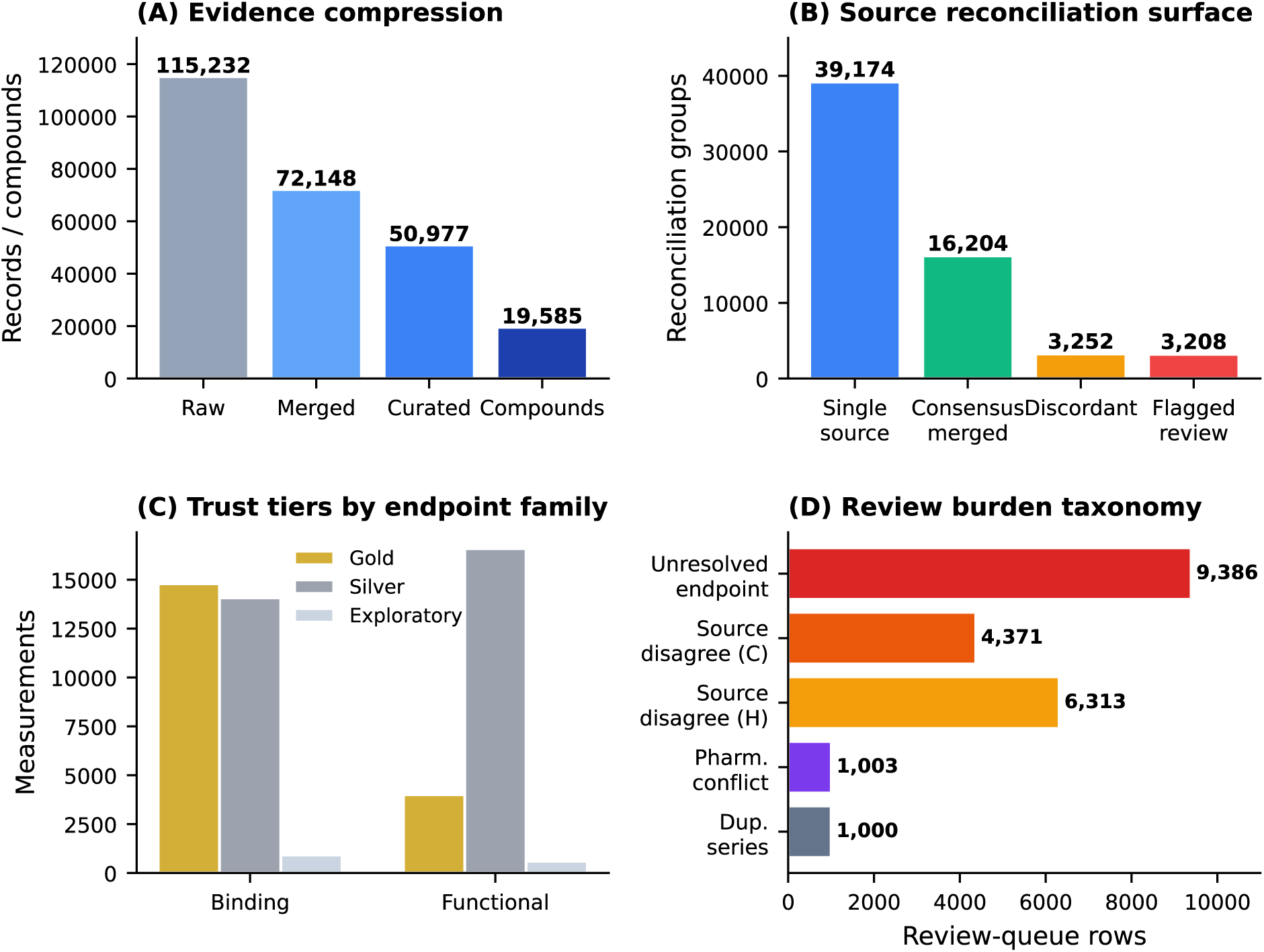
Curation converts heterogeneous public records into auditable evidence. (A) Evidence compression with audit sidecars: raw records, merged evidence, curated measurements, final compounds, explicit exclusions, review-queue records, and decision-lineage records. (B) Source-reconciliation surface: single-source, consensus-merged, discordant, and flagged-review groups. (C) Trust-tier structure by endpoint family. (D) Review-burden taxonomy showing the dominant uncertainty categories. Curation changes the evidentiary structure of the dataset rather than simply reducing row count.

Source reconciliation provides the first audit layer. Most reconciled groups were single-source (39,174) or consensus-supported (16,204), while 3,252 groups were discordant and 3,208 were flagged for review. These discordant and review-needed groups represent a non-negligible conflict surface in the public evidence base that, without explicit reconciliation, could be averaged, duplicated, or treated as independent support. By preserving reconciliation status, the workflow separates evidence breadth from evidence reliability. Full per-source reconciliation counts appear in Table S1.

Trust-tier assignment provides the second audit layer. Across the curated set, 18,797 measurements were classified as gold, 30,662 as silver, and 1,518 as exploratory. Within the binding family, 14,790 rows were gold, 14,070 silver, and 920 exploratory; within the functional family, 4,007 were gold, 16,592 silver, and 598 exploratory. The high retained gold/silver fraction shows that curation did not discard most public data; it preserved a large evidence base while making evidence quality explicit. The dominant review categories were unresolved endpoint meaning (9,386 development rows) and source disagreement (critical and high severity combined), followed by suspicious duplicate series, pharmacology conflict, duplicate activity inconsistency, and low-confidence assay semantics. Much of this uncertainty is semantic rather than numerical, concerning what was measured and how an assay should be interpreted. Full review-category counts by priority appear in Table S5.

The decision-lineage output is central to reproducibility: it records whether each record was retained, transformed, excluded, or flagged, allowing the benchmark to be audited at the level of curation decisions rather than only at the level of final metrics. The main conclusion from Figure 2 is that curation changes the evidentiary status of the benchmark. Raw records provide scale; reconciliation identifies agreement and conflict; standardization defines chemical identity; review queues identify unresolved ambiguity; trust tiers define benchmark-ready evidence; and decision lineage records how the dataset was created.

### Pharmacology- and readout-aware stratification defines functional QSAR tasks

After reconciliation and trust scoring, a distinct question remains: do the retained measurements define biologically coherent modeling tasks? This matters most for functional data. Binding measurements can often be interpreted as receptor-level affinity or displacement tasks, but functional measurements encode receptor subtype, endpoint scale, pharmacology class, assay readout, and signaling context. Treating all functional records as one pooled endpoint increases apparent sample size while weakening the biological meaning of the resulting claim. The endpoint ontology used to resolve functional records into benchmarkable tasks is shown in Figure 3. Functional evidence was stratified by assay readout and pharmacology class before matrix construction. Across the functional development pool, the dominant pharmacology classes were agonist (13,316 records), antagonist (2,704), and inhibitor (1,579), and the dominant assay readouts were binding displacement (33,017), GPCR generic (7,599), cAMP (1,501), and β- arrestin (520), with the remainder distributed across calcium, GTPγS, and generic functional readouts. The full readout-by-pharmacology coverage matrix appears in Table S6.

**Figure 3.**
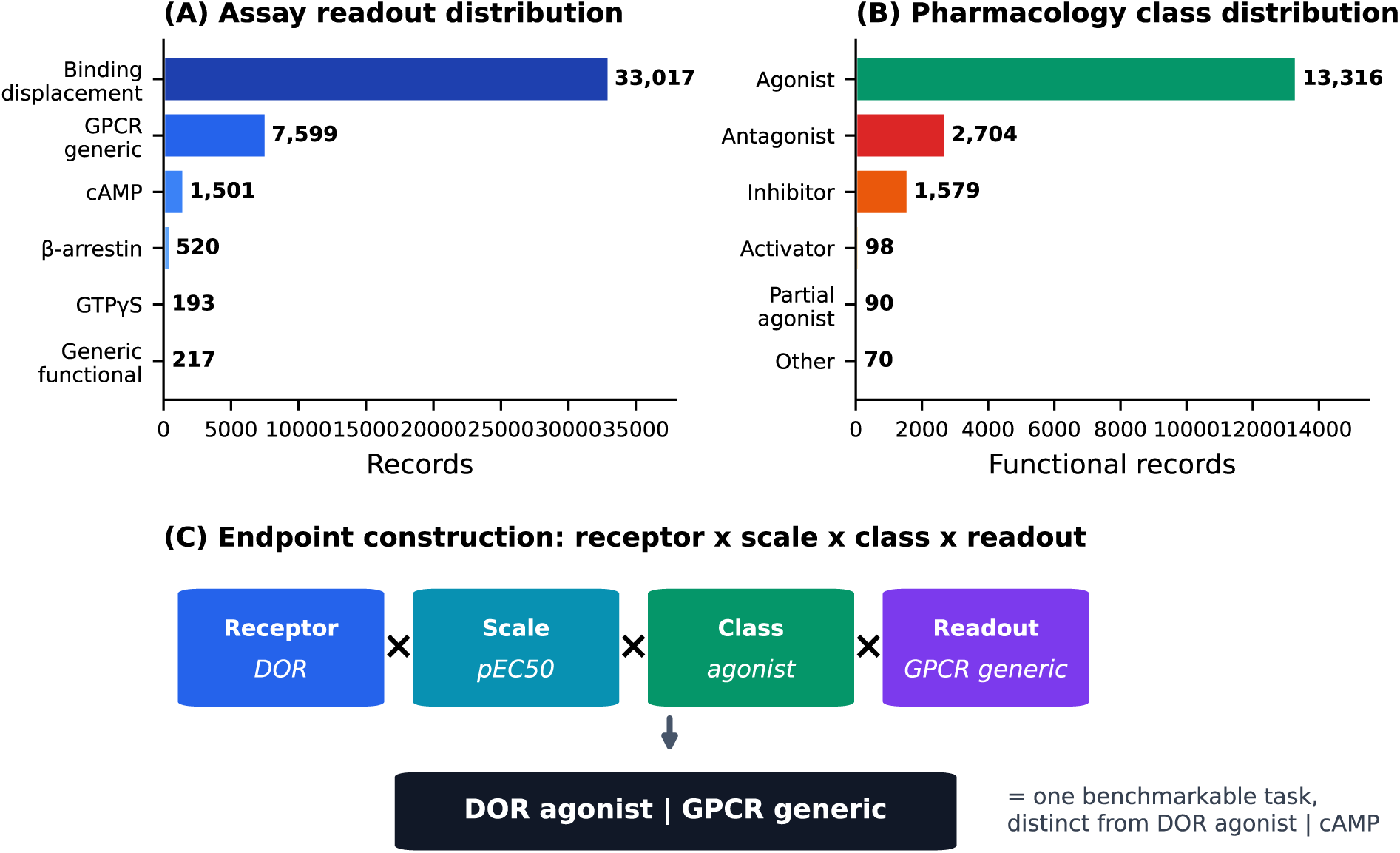
Endpoint ontology and pharmacology/readout stratification. (A) Functional evidence is many-to-many rather than one pooled endpoint, distributed across binding displacement, GPCR generic, cAMP, β-arrestin, generic functional, and other readouts, and across agonist, antagonist, and inhibitor pharmacology classes. (B) Endpoint construction rule showing how receptor, endpoint scale, pharmacology class, and assay readout define a benchmarkable functional task. (C) Examples of action/readout-resolved functional strata. This ontology prevents biologically distinct measurements from being collapsed into one ambiguous functional QSAR endpoint.

This separation is pharmacologically necessary.^(27)^ A cAMP assay measures Gs/Gi-mediated regulation of adenylyl cyclase; for the Gi/o-coupled opioid receptors, agonist activity often appears as inhibition of stimulated cAMP.^(33)^ A GTPγS or GPCR-generic activation assay measures G-protein activation nearer the receptor;^(34)^ a β-arrestin assay measures receptor-arrestin engagement relevant to biased signaling;^(35)^ and a binding-displacement readout measures radioligand competition and is therefore closer to an affinity measurement even when stored near functional labels.^(24)^ These readouts are not interchangeable views of one endpoint.^(23)^ A benchmarkable functional task is therefore defined by the combination of receptor, endpoint scale, pharmacology class, and assay readout: DOR + pEC_50_ + agonist + GPCR generic defines a task distinct from DOR agonist | cAMP, KOR antagonist | binding displacement, or MOR agonist | binding displacement. The endpoint name lengthens, but the claim sharpens.

This ontology also prevents binding-sentinel contamination. Some records filed under functional endpoint labels in fact describe radioligand binding or displacement; left in pooled functional models, they inject affinity information into functional potency tasks without pharmacological justification. By identifying binding-displacement readouts and routing them to their appropriate context, the workflow prevents functional models from training on mislabeled affinity-like measurements. The resulting benchmark includes multiple interpretable functional strata spanning agonist, antagonist, and inhibitor classes across distinct readouts, which a pooled functional label would collapse into a single ambiguous endpoint. The construction of the endpoint is therefore not discovered by the model; it is defined before modeling through pharmacology and readout resolution.

### External validation is meaningful only when interpreted with endpoint support

Once endpoint tasks are defined, performance can be read as a credibility landscape rather than a leaderboard (Figure 4). Locked external R^2^ is plotted against external sample support, annotated by endpoint identity, receptor, and class, so that a single performance number is not overinterpreted. The complete external-R^2^ landscape across all 71 externally evaluated endpoints (8 binding and 63 functional across the three functional arms) is reported per endpoint in Table S9, which also lists the internal-to-external generalization gap.

**Figure 4.**
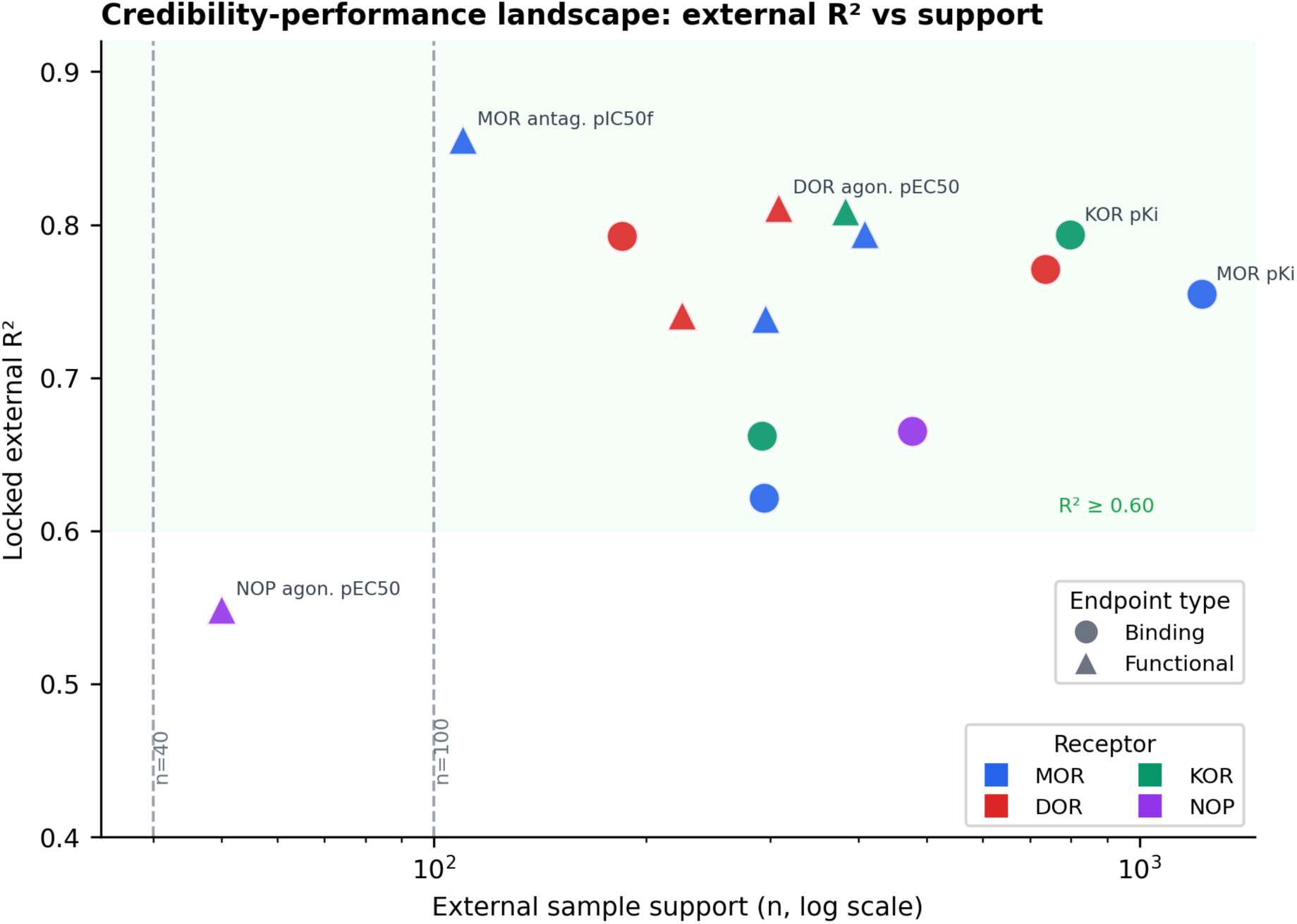
Credibility-performance landscape for externally validated opioid ligand QSAR endpoints. Locked external R² is interpreted together with endpoint identity and external sample support. Points represent binding and functional endpoints, colored by receptor and shaped by endpoint type. Support thresholds at n = 40 and n = 100 distinguish small, moderate, and broad-support claims. High external R² with sufficient support indicates a stronger claim, whereas high R² with small n is treated as hypothesis-generating. Model credibility depends on endpoint definition and evidence support, not R² alone.

The binding benchmark is the most compact and mature portion. The strongest receptor-level binding endpoint was KOR p*K_i_* (external R^2^ = 0.7937, consensus, n = 798), with DOR p*K_i_* (0.7712, LightGBM, n = 736), MOR p*K_i_* (0.7548, consensus, n = 1,227), and DOR pIC_50_ (0.7924, XGBoost, n = 185) also occupying the high-support, strong-performance region. NOP p*K_i_* reached R^2^ = 0.6650 (XGBoost, n = 477); after correction of the NOP target identity, this endpoint is a usable receptor-level binding result rather than the previously underdetermined task, and it is interpreted on its own footing rather than generalized from MOR, DOR, or KOR.

The functional benchmark is broader and more heterogeneous, and it is where stratification has the clearest effect. Action- and readout-resolved endpoints produced externally validated strata that a pooled functional label would not expose. The strongest functional result was a MOR antagonist functional-inhibition endpoint (R^2^ = 0.8562, consensus, n = 110). Action-stratified agonist potency endpoints were strong across receptors, including DOR agonist pEC_50_ (0.8114, XGBoost, n = 308), KOR agonist pEC_50_ (0.8094, LightGBM, n = 383), and MOR agonist pEC_50_ (0.7945, LightGBM, n = 408). When the same agonist potency was instead modeled through the broader GPCR-generic mechanistic stratum, performance remained solid (for example, DOR agonist | GPCR generic, R^2^ = 0.7415, consensus, n = 225; MOR agonist | GPCR generic, 0.7390, LightGBM, n = 295), illustrating that the action-stratified endpoints carry the stronger, more sharply defined claim.

The point is not that every functional endpoint is uniformly strong; it is that semantic resolution produces a more diverse panel of externally validated tasks. Of the 63 functional endpoints carried to external validation, 25 reached R^2^ of at least 0.60, and 23 of those combined R^2^ of at least 0.60 with external support of at least 40 compounds, meeting successive credibility thresholds. Because a high R^2^ on a small endpoint is not equivalent to the same value on a well-supported one, Figure 4 places endpoints in emerging, moderate, strong, and excellent regions using support bands at n = 40 and n = 100 rather than ranking by R^2^ alone. Bootstrap 95% confidence intervals for the top endpoints and the distribution-shift diagnostic (maximum mean discrepancy between training and external sets) are tabulated per endpoint in Table S9.

Model-family comparisons are intentionally not the main result. The strongest endpoints use different families, including LightGBM, XGBoost, and consensus models. This variation argues against a single-algorithm narrative: the dominant determinant of credibility is whether an endpoint is biologically defined, externally supported, and evaluated under scaffold-aware validation, not which model wins a leaderboard. Figure 4 therefore represents the endpoint-level consequence of the dataset contract, distinguishing mature, promising, and exploratory claims. Where a stratum remains small, such as the NOP functional endpoints, its external signal is treated as a direction for further data collection rather than a settled result, and it is reported on that footing in Table S9.

### Stratification is a precondition for a valid claim, not a performance optimization

The preceding results show that pharmacologically defined endpoints can be modeled with strong external performance. A separate and more fundamental question is what a model trained on pooled, unstratified functional data can actually claim. To test this directly, a fully pooled functional baseline was constructed for each receptor by combining all pharmacology classes and all assay readouts into a single endpoint per activity scale, with binding-sentinel rerouting disabled, and was trained and externally validated under the same scaffold-locked protocol as the stratified models. The comparison appears in Figure 5 and Table S10.

**Figure 5.**
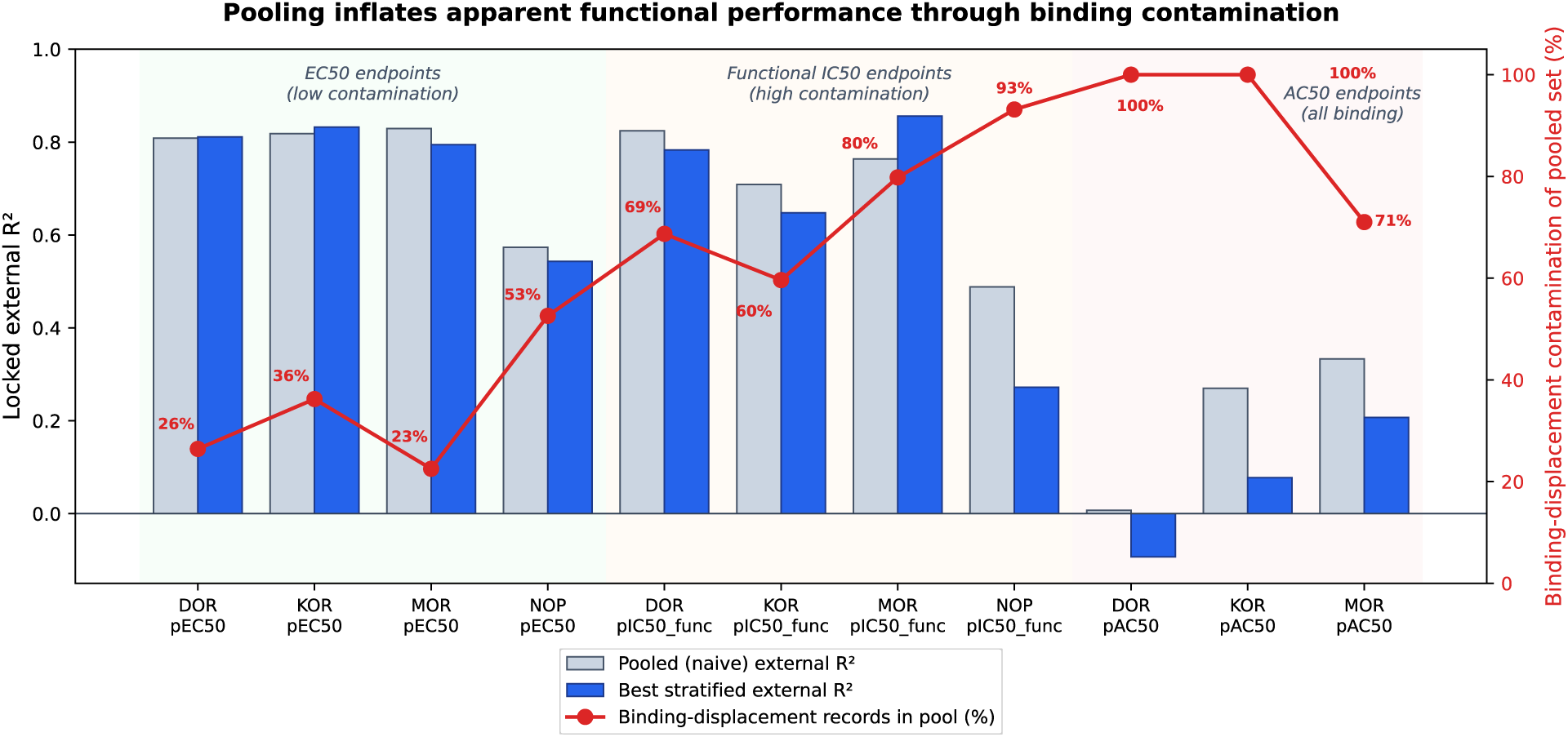
Pooling inflates apparent functional performance through binding-sentinel contamination. For each receptor and activity scale, locked external R² is shown for a fully pooled functional baseline (grey) and for the best pharmacology- and readout-stratified endpoint (blue), with the proportion of binding-displacement records in the pooled set overlaid (red line). For EC50 endpoints, where contamination is low, pooled and stratified models perform almost identically. For functional-IC50 and AC50 endpoints, where 60 to 100 percent of pooled records are binding-displacement measurements, the pooled model attains apparently strong R² by predicting binding affinity rather than functional potency. High pooled performance coincides with high contamination, illustrating that a pooled functional number measures a different quantity than its label claims.

The result is not that stratification uniformly raises R^2^. For agonist-dominated EC_50_ endpoints, where the pooled set is already pharmacologically coherent and contains little binding-displacement material (between 23 and 53 percent across receptors), pooled and stratified models performed almost identically (for example MOR pEC_50_, pooled R^2^ = 0.829 versus stratified 0.795; DOR pEC_50_, 0.809 versus 0.811; KOR pEC_50_, 0.818 versus 0.832). This parity is itself informative: where records already describe one pharmacology class in compatible readouts, stratification changes little, because there is little heterogeneity to resolve. The benefit of stratification appears precisely where heterogeneity is present, not as a uniform numerical gain.

The functional-IC_50_ endpoints expose why a pooled number can be high yet uninterpretable. Here the pooled set is dominated by binding-displacement records (between 60 and 93 percent), because radioligand-competition measurements filed under functional labels are no longer rerouted. The pooled DOR functional-IC_50_ model reaches R^2^ = 0.825, but 69 percent of its training records are binding-displacement measurements, so the model predicts binding affinity rather than functional inhibition; its apparent performance is borrowed from a different biological quantity than the endpoint label claims. The same pattern holds for KOR (R^2^ = 0.709, 60 percent binding) and MOR (R^2^ = 0.764, 80 percent binding). The AC_50_ pools are entirely binding-displacement (100 percent for DOR and KOR), and once isolated from any genuine functional signal they collapse (DOR pooled R^2^ = 0.007), confirming that the inflated functional-IC_50_ performance is contamination rather than predictive skill. Notably, for MOR the stratified antagonist functional-inhibition endpoint (R^2^ = 0.8562, n = 110) outperforms the contaminated pool (0.764) even though the pool is 80 percent binding, because isolating the genuine functional signal yields a stronger, coherent task.

The deeper issue is that a pooled functional model cannot support a defensible scientific claim, independent of its R^2^. A single model trained across agonist and antagonist records and across cAMP, [^35^S]GTPγS, β-arrestin, and binding-displacement readouts cannot answer the question it would need to answer, namely what activity it predicts, because there is no single functional activity that these assays jointly measure. The same agonist can be potent in one functional readout and markedly weaker in another as a consequence of signaling bias and assay coupling rather than experimental error, so a model regressing structure against the pooled value learns a quantity that corresponds to no real assay outcome. Its prediction cannot tell a medicinal chemist whether a scaffold will be a potent G-protein agonist or an arrestin recruiter, which is exactly the distinction that pathway-selective opioid design depends on. A stratified endpoint, by contrast, supports a testable statement, that a model predicts agonist potency in a defined readout, whereas a pooled endpoint supports only an uninterpretable average. Stratification by pharmacology class and assay readout is therefore a precondition for the model output to constitute a valid claim, not an optimization applied to improve a metric.

This readout dependence is observable directly in the curated data, although the relevant overlap is uncommon because most studies report a single functional assay per compound. In one systematic series of opioid receptor agonists, the same compounds were characterized both by Gα-coupled activation (EC_50_) and by inhibition of forskolin-stimulated cAMP (IC_50_); across fifteen compounds measured by both readouts, apparent agonist potency differed by a median of roughly 0.9 log units and by up to approximately 2 log units between the two formats.^(36)^ Because such cross-readout measurement of the same compound is rare, a pooled functional model seldom observes these differences directly and instead averages across endpoint definitions and assay formats that are not interchangeable, which is precisely why the pooled value cannot be tied to a single, well-defined pharmacological claim.

### SHAP attribution separates binding and functional structure-activity signals

The benchmark includes a SHAP interpretation layer to examine whether curated binding and functional tasks encode different descriptor-level signals. SHAP was used as an interpretability layer rather than as proof of mechanism, and the analysis was anchored to Random Forest models so that descriptor contributions could be compared consistently across endpoints and splits, even where another family produced the best external R^2^. Across the two benchmark modes, the workflow produced 100 Random Forest SHAP summary panels, comprising 8 binding panels and 92 functional panels across the naive, balanced, and maximal functional arms. Because scaffold transfer is the stricter generalization setting, the main-text interpretation focuses on scaffold models; the complete panel set is released as Supporting Information and summarized in Section S4.4.

The binding panels cover receptor-level MOR, DOR, KOR, and NOP models under the scaffold split and support the view that binding tasks are comparatively compact but still differ by receptor in both performance and descriptor attribution. For the Random Forest models that anchor the SHAP analysis, locked external performance on receptor p*K_i_* was 0.7744 for KOR (n = 798), 0.7558 for DOR (n = 736), 0.7445 for MOR (n = 1,227), and 0.5905 for NOP (n = 477), so the binding interpretation rests on models with substantial external support across the family. The functional panels cover action/readout-resolved tasks aligned with the main claims, spanning broad-support agonist endpoints and smaller antagonist and inhibitor readouts.

The biological interpretation is that binding and functional models do not rely on identical descriptor patterns (Figure 6). Binding models tend to emphasize scaffold identity, local substructure environments, and receptor-specific affinity determinants. Functional models, especially agonist and antagonist tasks, place greater weight on descriptors related to nitrogen charge environment, molecular topology, branching, hydrophobic distribution, and polarizable surface area. These descriptors are chemically meaningful for opioid ligands: the basic nitrogen is central to receptor binding and receptor-state stabilization, while topology and hydrophobic distribution influence how a ligand occupies the orthosteric pocket and whether it supports a functional receptor state. Functional potency is not simply binding affinity plus noise; a ligand can bind strongly yet differ in agonist efficacy, antagonist behavior, β-arrestin recruitment, or G-protein activation. The SHAP layer is consistent with this distinction and reinforces the curation argument: if binding and functional potency encode different signals, pooling binding-like and functional records, or pooling functional readouts without pharmacological resolution, risks hiding the very differences medicinal chemistry needs to exploit. These patterns are presented as descriptor-level signals consistent with known opioid SAR, not as causal proof of receptor mechanism.

**Figure 6.**
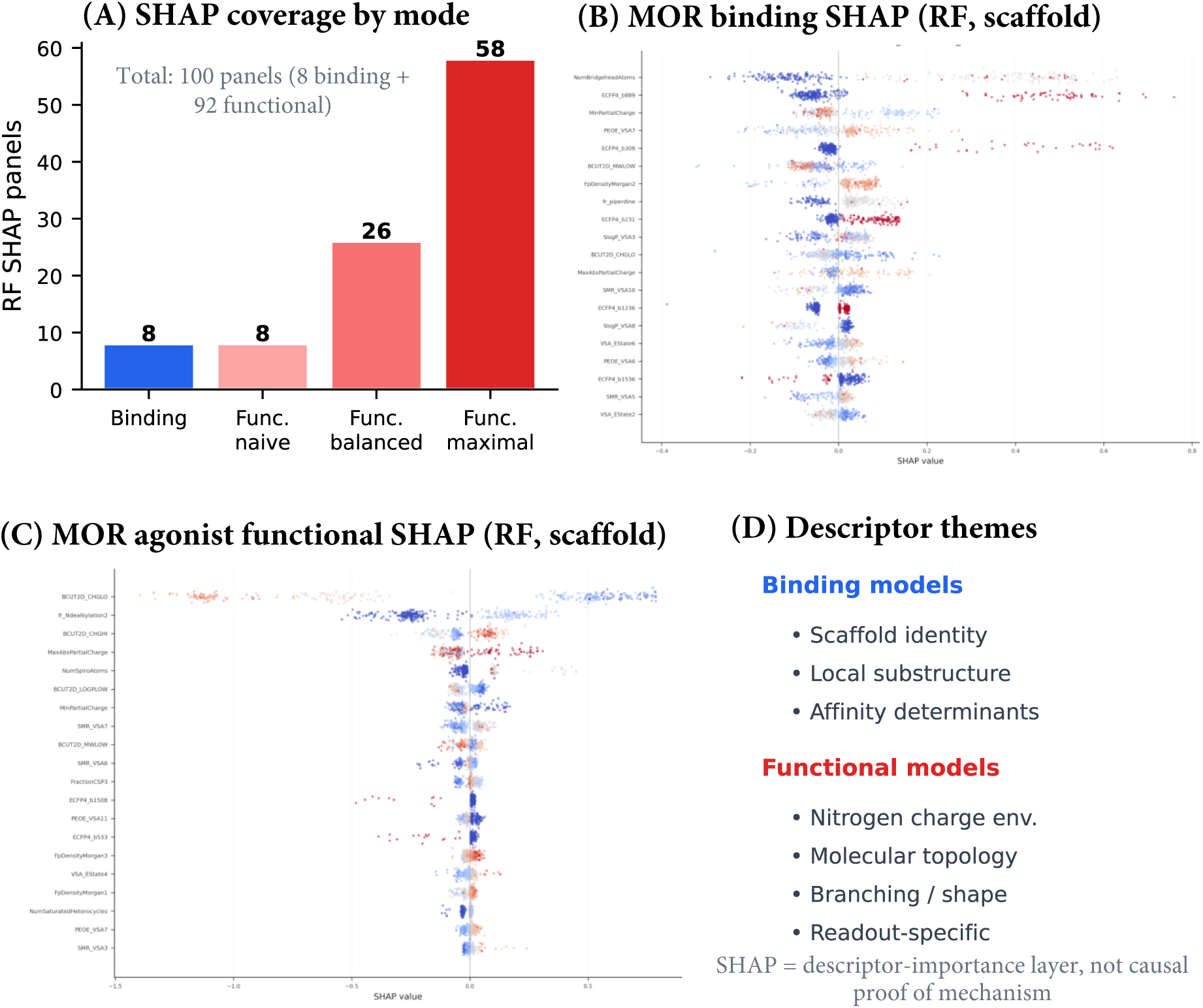
SHAP interpretation layer for scaffold-aware opioid ligand QSAR. (A) Coverage of Random Forest SHAP exports by benchmark mode and split, with scaffold panels emphasized because scaffold transfer is the stricter generalization setting. (B) Selected scaffold SHAP endpoints used for the main-text interpretation, showing external R² and support for receptor-level binding and action/readout-resolved functional tasks; small-n endpoints are marked and interpreted cautiously. (C) Descriptor themes: binding models emphasize scaffold and local substructure identity, whereas functional models emphasize nitrogen charge environment, molecular topology, branching, conformational shape, and action/readout-specific signals. SHAP is used as a descriptor-importance layer, not as causal proof of mechanism.

## Discussion

### Data curation as the primary determinant of QSAR credibility

The central conclusion of this study is general and not specific to one target family: the validity of a QSAR claim is determined first by data curation, not by model architecture, and validity is not the same as predictive performance. The fully pooled baseline makes this concrete. On agonist-dominated EC_50_ endpoints, which are already coherent, pooled and stratified models perform almost identically, so a high R^2^ on a pooled model is not by itself evidence that pooling is sound. On functional-IC_50_ endpoints, the pooled model can reach a strong R^2^ while the majority of its records are binding-displacement measurements, so the number is high precisely because the model has quietly become a binding-affinity predictor. Performance therefore cannot certify a claim; only the dataset definition can. A model trained on a poorly defined endpoint can emit accurate-looking predictions without supporting any clear pharmacological statement, whereas a model trained on a smaller but pharmacologically coherent endpoint supports a claim that is testable and falsifiable. In the opioid demonstration, readout-aware functional stratification produced externally validated action/readout-resolved strata, including action-stratified agonist potency endpoints for DOR, KOR, and MOR and a MOR antagonist functional-inhibition endpoint that outperforms the contaminated pooled alternative; the value of stratification is that each of these endpoints corresponds to a real, single assay outcome that a medicinal chemist can act on. Model choice still matters but is secondary: different families won on different endpoints, which supports per-endpoint reporting rather than a single model-family comparison.

### Assay-readout stratification as pharmacological rigor

The principal pharmacological contribution of the framework is the separation of functional endpoints by action type and assay readout. Opioid functional assays are not interchangeable measurements of one quantity: cAMP, GTPγS, β-arrestin, and binding-displacement readouts report different events, and treating them as a single functional endpoint can obscure the biology that opioid pharmacology aims to understand. This is especially relevant to biased agonism,^(37)^ which depends on differences between pathways such as G-protein activation versus β-arrestin recruitment. A model trained on pooled readouts cannot resolve bias because it averages over the pathway differences that bias exploits, whereas a readout-resolved model can ask whether structure predicts potency in a specific receptor-pharmacology-readout context.^(13)^ The binding-displacement strata deserve particular attention: rerouting radioligand-competition records away from functional labels prevents binding-like measurements from contaminating functional potency models, which changes the scientific question being modeled rather than serving as a cosmetic cleanup. The pooled baseline quantifies the cost of skipping this step, since functional-IC50 pools that retain 60 to 100 percent binding-displacement records produce models that predict affinity under a functional label. For heterogeneous GPCR data more broadly, endpoint family, endpoint scale, receptor identity, pharmacology class, and assay readout should be treated as separate components of the endpoint definition; the more heterogeneous the assay biology, the more hazardous a single endpoint label becomes.

### Generality and integration with existing curation resources

Although demonstrated here in depth on four opioid receptors, the framework is constructed as a general curation methodology rather than an opioid-specific pipeline. Its operative primitives, namely source reconciliation with preserved agreement metadata, deterministic chemical standardization, pharmacological action-type assignment, binding-sentinel detection, assay-readout stratification, trust-tier scoring, scaffold-locked partitioning, and decision-lineage recording, are defined in terms of pharmacology and assay semantics that apply to any receptor or enzyme target class, not in terms of opioid-specific chemistry. The pharmacological knowledge that drives stratification is supplied as an external, versioned configuration rather than embedded in the analysis code, so that extending the approach to a new target class is a matter of describing its assays and action types rather than rewriting the workflow.

The approach is also designed to compose with, rather than replace, existing standardization efforts. Large-scale resources that harmonize structures, units, and activity types^(20)^ address the measurement-type and chemical-representation axes of heterogeneity; the pharmacological-stratification axis introduced here is orthogonal to them and can be layered on top of such standardized collections. In practice, a practitioner could take a standardized bioactivity set, apply the action-type and readout stratification described in this study, and obtain pharmacologically coherent modeling tasks without discarding the upstream standardization. The contribution is therefore additive to the existing data-curation ecosystem: it supplies the semantic layer that converts standardized-but-pooled records into endpoints whose biological meaning is explicit.

### SHAP interpretation and implications for ligand design

The SHAP layer adds a descriptor-level view of the curated benchmark. It is anchored to Random Forest models for a consistent attribution lens across receptors, endpoint families, and splits, which should not be read as a claim that Random Forest is the best family for every task. Binding panels indicate substantial external support and comparatively stable scaffold-transfer behavior, with MOR, DOR, and KOR showing stronger transfer than NOP, a difference best interpreted as endpoint-specific learnability rather than failure of the binding benchmark. Functional panels span broad-support agonist endpoints and smaller antagonist and inhibitor readouts, reinforcing the credibility principle that strong descriptor patterns in small-n endpoints are hypothesis-generating while broad-support endpoints support more mature interpretation. The descriptor themes are chemically plausible for opioid ligands: binding emphasizes scaffold and local substructure identity, while function emphasizes nitrogen electronics, topology, branching, conformational shape, and readout-specific signals. These are interpretive hypotheses rather than causal mechanisms, but they link the curated endpoint structure to testable SAR ideas, for example that modifying an N-substituent may alter both the protonated-nitrogen environment and molecular topology and thereby change functional potency even when binding remains high.

### OECD-aligned, dataset-contract reporting

The framework aligns naturally with OECD-style QSAR reporting because it makes endpoint definition, algorithm, applicability domain, validation, and interpretation explicit,^(38)^ but it treats these not as a closing checklist. They are built into the dataset contract. Endpoint definition occurs before training; external validation is locked before performance is interpreted; evidence quality is represented through trust tiers and review queues; source conflict is preserved through reconciliation metadata; and SHAP is added after validation as an interpretability layer rather than a substitute for it. This ordering prevents model performance from being interpreted independently of the curation decisions that created the endpoint, and it makes the released artifacts part of the scientific method rather than supplementary material.

### Outlook

Two directions follow naturally from this work and define where the approach extends next. First, the breadth of pharmacological resolution can be pushed further: the maximal-resolution functional arm shows that as endpoints are subdivided into increasingly specific pharmacology-readout strata, some strata become small enough that external estimates on them are dominated by a few high-variance points and are best summarized by support-aware, endpoint-level statistics rather than by pooled averages. This behavior is itself informative, marking the frontier at which additional data, rather than additional resolution, becomes the limiting factor, and the trust-tier and review-queue machinery is designed to flag exactly these strata for targeted data collection. The NOP functional endpoints are the clearest current example: NOP binding is now a well-supported receptor-level task, while NOP functional potency is best treated as an emerging, hypothesis-generating direction for which the framework already identifies the specific strata that would benefit from new measurements. Second, because the pharmacological knowledge driving stratification is externalized as a versioned configuration, the same framework can be carried to other receptor and enzyme families and composed with existing standardization resources, extending pharmacology-aware curation across the broader bioactivity ecosystem.

### Overall interpretation

The main result of this study is not a single best model. It is a framework for deciding which QSAR claims are credible, demonstrated where the stakes are high. Public bioactivity data contain useful signal, but that signal becomes scientifically interpretable only when the dataset defines source provenance, chemical identity, endpoint meaning, pharmacology class, assay readout, evidence quality, validation population, and interpretability context. The model does not define the claim alone; the curated endpoint does.

## Materials and Methods

### Target selection and four-source public data retrieval

Bioactivity records were curated for four human opioid receptor targets: the mu opioid receptor (MOR; CHEMBL233), delta opioid receptor (DOR; CHEMBL236), kappa opioid receptor (KOR; CHEMBL237), and nociceptin/orphanin FQ receptor (NOP/NOR; CHEMBL2014). At first mention the nociceptin receptor is denoted NOP/NOR to acknowledge both naming conventions; NOP is used thereafter for consistency with the figures and tables.^(39)^ A target-scoped retrieval strategy was applied across four resources: ChEMBL, BindingDB, PubChem, and GtoPdb, with the public databases accessed in June, 2026 and the retrieved snapshots frozen for reproducibility. Records were retained when mapped to one of the four targets and containing sufficient compound, structure, activity, endpoint, and source metadata for curation or review. Retrieved fields included source name, source compound identifier, target identifier and name, endpoint label, numerical value, unit, activity relation operator, assay identifier, assay description, assay type, document identifier where available, source record identifier, compound structure, and source-provided action or mechanism annotation where available. Retrieval produced 115,232 raw source rows, harmonized into a common schema and merged into 72,148 records before downstream preparation.

### Source normalization and reconciliation

Source-specific records were normalized before merging. Target identifiers were mapped to MOR, DOR, KOR, or NOP; endpoint labels were normalized to canonical names such as K*_i_*, K*_d_*, IC_50_, EC_50_, AC_50_, Emax, efficacy, and inhibition; units were standardized; activity relation operators were mapped to a common vocabulary; and structures were passed forward for deterministic standardization. Records were reconciled by compatible compound identity, target, endpoint family, activity scale, and source provenance, with evidence-group categories assigned as single-source, consensus-merged, discordant, or flagged for review. Multi-source overlap was treated as reliable only when source annotations and values were sufficiently concordant; discordance and review-needed status were preserved as dataset attributes and propagated into trust-tier scoring and decision-lineage outputs.

### Pre-standardization structural filters

Four structural quality filters were applied sequentially before standardization. First, each SMILES string was parsed with MolFromSmiles followed by SanitizeMol;^(40)^ records that failed were excluded and logged. Second, disconnected mixtures were detected by fragment enumeration with GetMolFrags and excluded when two or more disconnected organic fragments were present, because identity and descriptor calculation become ambiguous for multi-component mixtures.^(41)^ Third, inorganic compounds (zero carbon atoms after explicit-hydrogen handling) were removed. Fourth, organometallic structures with direct metal-carbon bonds for metallic elements from lithium through americium were removed, because standard small-molecule descriptor assumptions are less reliable for them. All exclusions were written to the compound-level curation decision log with the corresponding reason.

### Chemistry standardization

Validated structures underwent a deterministic seven-step cascade in fixed order: (1) salt stripping and largest-fragment selection, relevant for opioid ligands frequently deposited as hydrochloride, sulfate, or tartrate salts; (2) bond and charge normalization for nitro groups, charge-separated structures, and protonation variants; (3) charge neutralization to neutral parent structures where chemically appropriate, without forcing zwitterionic or permanently charged species into invalid forms; (4) explicit-hydrogen removal with RemoveHs; (5) tautomer canonicalization using TautomerEnumerator followed by Canonicalize(), relevant for keto-enol, amide-iminol, and heteroatom-rich motifs; (6) canonical SMILES generation in isomeric and non-isomeric forms; and (7) identifier generation with stereochemistry-aware and connectivity-only InChIKey-style identifiers. This cascade produced standardized structures, canonical strings, and normalized identifiers used for deduplication, scaffold assignment, descriptor generation, and leakage audits. Across the curated set, 70,947 input structures were processed and valid, of which 14,445 were changed by standardization, 1,944 were fragment-collapsed or had salts and solvents removed, and 12,423 were tautomer-canonicalized; 1,537 carried PAINS alerts and 37 were isotope-labeled inputs.

### Physicochemical property and PAINS annotation

For each standardized compound, physicochemical properties were computed to support quality control and chemical-space interpretation, including molecular weight, calculated LogP, topological polar surface area, hydrogen-bond donor and acceptor counts, rotatable-bond count, ring and aromatic-ring counts, heavy-atom count, fraction of sp^3^ carbon, and quantitative estimate of drug-likeness.^(42)^ PAINS screening used substructure filters for common assay-interference motifs;^(43)^ flagged compounds were excluded from primary training sets unless otherwise specified for sensitivity analysis, with drug-like annotations retained as metadata.

### Activity standardization and censored-data handling

Activity values were converted to a common pActivity scale when the endpoint represented a molar concentration, pActivity = −log_10_(value × unit factor), with unit factors nM = 10^−9^, μM = 10^−6^, mM = 10^−3^, and M = 1. Values already on a logarithmic scale (pK*_i_*, pIC_50_, pEC_50_, pK*_d_*) were retained directly, and percentage endpoints such as inhibition, activity, Emax, and efficacy were kept on their native scale. Activity relation operators were preserved as censored-data annotations: greater-than values as upper bounds and less-than values as lower bounds on the pActivity scale. Primary regression models and metrics were computed on exact measurements only; censored values were retained in curated tables for audit and future censored-model extensions.

### Endpoint family, action-type annotation, and binding-sentinel detection

Each record was assigned an endpoint family before modeling. Binding endpoints included K*_i_*, K*_d_*, pK*_i_*, pK*_d_*, binding IC_50_, and records whose assay descriptions indicated radioligand binding or displacement. Functional endpoints included EC_50_, pEC_50_, AC_50_, Emax, efficacy, functional IC_50_, and inhibition of a functional response. Endpoint labels were treated as provisional until assay semantics were assigned, because IC_50_ can represent binding displacement, functional inhibition, or antagonist blockade. Functional records were assigned pharmacological action types through a priority cascade: explicit source-provided annotation first, then assay-description text mining, then endpoint and readout context, covering agonist, partial agonist, inverse agonist, antagonist, inhibitor, blocker, allosteric modulator, activator, binding, and unresolved classes. A binding-sentinel step detected records filed under functional labels but describing radioligand binding or displacement (for example, competition binding or displacement of labeled ligand) and rerouted them to binding-displacement contexts, preventing affinity measurements from contaminating functional potency models.

### Assay-readout stratification and endpoint construction

Functional records were stratified by assay readout because different readouts measure different processes. Readout categories included GPCR cAMP, GPCR generic/GTPγS-like activation, β-arrestin recruitment, binding displacement, calcium mobilization, generic functional response, and other low-frequency or unresolved readouts. Endpoint columns were then constructed to include receptor, endpoint scale, pharmacology class, and assay readout, so that a DOR pEC_50_ agonist record in a GPCR-generic assay was assigned to a DOR agonist | GPCR generic task. This prevented agonist, antagonist, inhibitor, and readout-specific records from being pooled into one ambiguous functional endpoint. To probe the effect of stratification depth, functional endpoints were constructed at three resolutions, a naive arm with minimal stratification, a balanced arm, and a maximal-resolution arm that subdivides into the most specific pharmacology-readout strata; all three are derived from the same curated long table.

### Deduplication, quality filtering, and trust-tier scoring

Replicate measurements were grouped by standardized identity, target, endpoint scale, endpoint family, pharmacology class, and assay readout. Exact-measurement groups used the median pActivity; upper-bound-only groups retained the minimum and lower-bound-only groups the maximum as conservative bounds; mixed-relation groups were retained with metadata and flagged when interpretation required review. Inter-replicate standard deviation above 1.0 log unit flagged activity inconsistencies. Quality filters retained compounds with molecular weight 100-900 Da, pActivity 2.0-12.0 for molar potency endpoints, and heavy-atom count 5-100; PAINS-flagged records were excluded from primary training; Lipinski and Veber criteria were retained as annotations. Each retained record received a row-level trust score combining source support, reconciliation status, assay-semantic confidence, relation status, replicate agreement, endpoint-family confidence, pharmacology-class confidence, and readout specificity, mapped to gold, silver, or exploratory tiers. A review queue captured records needing attention, with categories for unresolved endpoint meaning, source disagreement, pharmacology conflict, suspicious duplicate activity inconsistency, suspicious duplicate series, and low-confidence assay semantics. Records failing filters were not silently removed; each exclusion was written to the decision log.

### Scaffold computation, dataset splitting, and applicability domain

Bemis-Murcko scaffolds and generic scaffolds were computed from standardized structures.^(21)^ A scaffold-locked external holdout was reserved before model development, with scaffold families assigned exclusively to development or external partitions to reduce leakage; the external set comprised 7,780 measurements across 2,937 compounds (15.0% of compounds; seed 42), and the development set 43,197 measurements across 16,648 compounds, spanning 7,544 total scaffolds of which 1,179 were external. Within the development set, internal validation used both scaffold and random splits, with random splits as optimistic reference baselines and scaffold splits assigning entire scaffold groups to a single partition. Leakage was audited by Morgan-fingerprint Tanimoto similarity, flagging external compounds with maximum training similarity above 0.9. Applicability domain was assessed using leverage h*_i_* = x*_i_*ᵀ(XᵀX)⁻¹x*_i_*, with warning threshold h* = 3p/n for p features and n training compounds. Internal split summary statistics appear in Table S7.

### Molecular descriptors and model training

Two descriptor families were used: RDKit two-dimensional descriptors (constitutional, electronic, topological, surface-area, charge, fragment, and physicochemical) and Morgan circular fingerprints at radius 2 in an ECFP4-like representation.^41^ Descriptor matrices were cleaned by removing constant, all-missing, and infinite columns, imputing remaining missing values with zero; feature scaling was fit on the training set only and applied to test and external sets. Five model families were trained per endpoint and split: Random Forest (300 trees, square-root feature sampling, minimum leaf size 2), XGBoost (300 rounds, depth 6, learning rate 0.1, subsample 0.8, column subsample 0.8), LightGBM (300 rounds, leaf-wise growth, learning rate 0.1, subsample 0.8, column subsample 0.8), Support Vector Regression (RBF kernel, C = 10.0, ε = 0.1, scale-based γ), and a consensus model averaging Random Forest, XGBoost, and Support Vector Regression. Hyperparameter tuning, when enabled, used training data only; external data were never used for scaling, model selection, tuning, or threshold selection. The complete hyperparameter configuration appears in Table S1 and per-model leaderboard statistics in Table S8.

### Evaluation metrics and validation analyses

Performance was assessed using the coefficient of determination R^2^, root-mean-square error, and mean absolute error, computed on exact measurements only, where R^2^ = 1 − Σ(y*_i_* − ŷ*_i_*)² / Σ(y*_i_* − ȳ)². Six validation tiers were applied: (1) a scaffold-locked external set for the primary claim, interpreted with endpoint support and sample size; (2) 500-resample bootstrap confidence intervals where sample size allowed (reported in Table S9); (3) distribution shift quantified by maximum mean discrepancy with an RBF kernel and median-heuristic γ (reported in Table S9); (4) scaffold-aware GroupKFold cross-validation using generic scaffolds as group labels with five folds; (5) y-randomization with 20 label permutations compared against the permuted-label null; and (6) conformal prediction intervals at 90% and 95% where support was sufficient. Activity-cliff directional recall was computed for relevant endpoints as the fraction of cliff pairs whose predicted potency-difference sign matched observation.

### SHAP feature importance

SHAP feature attribution interpreted descriptor contributions for tree-based Random Forest models.^(38)^ SHAP values were computed for scaffold-split test compounds, with subsampling for large test sets, and global importance summarized as the mean absolute SHAP value across evaluated compounds. The main-text interpretation focuses on scaffold-split models because scaffold transfer is the stricter generalization setting. The SHAP export layer contained 100 Random Forest summary panels, comprising 8 binding and 92 functional panels across the naive, balanced, and maximal functional arms. The SHAP layer is used as a descriptor-importance and SAR-hypothesis layer, not as causal proof of receptor mechanism, and is anchored to Random Forest rather than necessarily to the best-performing family for each endpoint.

### OECD-aligned reporting

The benchmark was organized to align with OECD-style QSAR reporting. Principle 1, defined endpoint, is addressed by reporting each endpoint with receptor, endpoint scale, endpoint family, pharmacology class, and assay readout. Principle 2, unambiguous algorithm, is addressed through declared descriptors, model families, split logic, hyperparameters, and consensus definition. Principle 3, applicability domain, is addressed through scaffold splitting, fingerprint similarity, leverage analysis, and distribution-shift diagnostics. Principle 4, goodness-of-fit, robustness, and predictivity, is addressed through internal validation, locked external validation, bootstrap confidence intervals, y-randomization, and scaffold-aware cross-validation. Principle 5, mechanistic interpretation, is addressed through endpoint semantics, assay-readout biology, pharmacology class, and SHAP attribution.

## Conclusions

This study shows that the validity of a QSAR claim is determined first by how the dataset is defined, and that pharmacological mechanism and assay readout constitute a primary curation axis distinct from the measurement-type and chemical-representation axes addressed by existing standardization. The framework integrates multi-source public records into an auditable evidence structure recording source agreement and conflict, chemical identity, endpoint meaning, assay readout, pharmacology class, trust tier, review status, and decision lineage, and it stratifies functional data by action type and readout before modeling. Demonstrated on the four opioid receptors, it reconciled 115,232 raw records into 72,148 merged records, 50,977 curated measurements, and 19,585 compounds, and produced receptor-level binding benchmarks with strong locked external performance together with action/readout-resolved functional strata. A direct comparison against a fully pooled functional baseline shows why this stratification is necessary: pooled functional models either match the stratified models on already-coherent endpoints, offering no advantage, or attain a deceptively high R^2^ on functional-IC50 endpoints by training predominantly on binding-displacement records, so that the pooled number predicts affinity rather than the labeled functional activity. The stratification is therefore a precondition for a model output to constitute a defensible claim, because a model pooled across agonist and antagonist records and across distinct functional readouts predicts a quantity that corresponds to no single assay outcome, whereas a stratified endpoint supports a testable statement about activity in a defined pharmacological and readout context. SHAP attribution further indicates that binding and functional tasks encode partially distinct structure-activity signals. Because its pharmacological knowledge is externalized as a versioned configuration and its primitives are defined in assay-semantic terms, the framework is reusable across target classes and composes with existing standardization resources, supplying the semantic layer that turns standardized-but-pooled records into endpoints whose biological meaning is explicit.

## Data availability

The data and code supporting this article are available as Supporting Information and in a public repository, CURATO-OPIOID, at https://github.com/kelokely/CURATO-OPIOID, released under the Creative Commons Attribution-NonCommercial 4.0 (CC BY-NC 4.0) license. The repository provides the curated bioactivity datasets, a versioned pharmacology and assay-readout ontology that defines the stratification, a reference implementation of the model training and scaffold-locked external validation, and the fully pooled versus stratified comparison, sufficient to reproduce the reported endpoint-level results. The pharmacological knowledge that drives stratification is supplied as a separate, version-pinned configuration file rather than embedded in code, so that the curation logic is inspectable and extensible to other target classes. Molecular visualization and property review used MolScope (https://kelokely.github.io/-molprop-toolkit/).

## Supporting information

SI description

## Author contributions

K.M.E. conceptualized the study, designed the curation and benchmarking framework, and supervised the work. M.A.N. contributed to software. M.A.N. and L.M.A. contributed to methodology, and data curation. A.G., S.J.W., L.-Y.L.-C., A.M.R., and M.A.-G. contributed to pharmacological interpretation and validation. K.M.E. wrote the original draft; all authors reviewed and edited the manuscript. All authors have read and agreed to the submitted version.

## Acknowledgements

This publication was made possible by an Institutional Development Award (IDeA) from the National Institute of General Medical Sciences of the National Institutes of Health under Grant 2P20GM103432. The content is solely the responsibility of the authors and does not necessarily represent the official views of the National Institutes of Health. The authors thank the maintainers of ChEMBL, BindingDB, PubChem, the IUPHAR/BPS Guide to Pharmacology, and RDKit, and the broader scientific Python ecosystem for making public-data cheminformatics benchmarking possible.

