## Supplementary material for "Pharmacological Stratification of Public Bioactivity Databases: A Reusable, OECD-Anchored Curation and Benchmarking Framework Demonstrated for Opioid Receptors": SI

### Contents

This document contains supporting tables (Tables S1-S10) and a description of the supporting data. Table S1 lists model hyperparameters. Tables S2 and S3 describe source contributions and the source-by-target retrieval matrix. Tables S4-S6 describe trust tiers, review-queue structure, and the functional readout-by-pharmacology coverage. Tables S7 and S8 summarize internal split statistics and the per-model external leaderboard. Table S9 reports per-endpoint external validation with confidence intervals and distribution-shift diagnostics. Table S10 reports the fully pooled versus stratified functional comparison. Section S4.4 summarizes the SHAP analysis, and Section S5 lists the supporting data files.

**Table S1.** Hyperparameter values used for the five model families. Values were fixed across all endpoints and split regimes; hyperparameter tuning was not used so that performance differences could be attributed to dataset characteristics rather than endpoint-specific tuning.

| Model family | Key hyperparameters |
|---|---|
| Random Forest (RF) | 300 trees; max features = sqrt; min samples per leaf = 2; bootstrap = true |
| XGBoost (XGB) | 300 rounds; max depth = 6; learning rate = 0.1; subsample = 0.8; colsample_bytree = 0.8 |
| LightGBM (LGBM) | 300 rounds; leaf-wise growth; learning rate = 0.1; subsample = 0.8; colsample_bytree = 0.8 |
| Support Vector Regression (SVR) | RBF kernel; C = 10.0; epsilon = 0.1; gamma = scale |
| Consensus | Arithmetic mean of RF, XGB, and SVR predictions |

*Descriptor representation for all model families: RDKit two-dimensional descriptors and Morgan fingerprints at radius 2 (ECFP4-like). Feature scaling was fit on the training partition only.*

**Table S2.** Per-source contribution to the integrated opioid receptor bioactivity resource. Raw counts are records retrieved before merge; merged counts are records after source reconciliation.

| Source | Raw records | Merged records |
|---|---|---|
| PubChem | 46,841 | 30,179 |
| BindingDB | 36,184 | 15,309 |
| ChEMBL | 31,940 | 26,504 |
| IUPHAR/BPS Guide to Pharmacology | 267 | 156 |
| Total | 115,232 | 72,148 |

**Table S3.** Source-by-target retrieval matrix (raw records before merge and curation). Columns are the four opioid receptors; the nociceptin receptor is CHEMBL2014.

| Source | MOR (233) | DOR (236) | KOR (237) | NOP (2014) | Total |
|---|---|---|---|---|---|
| BindingDB | 12,911 | 8,131 | 11,485 | 3,657 | 36,184 |
| ChEMBL | 11,443 | 7,058 | 9,902 | 3,537 | 31,940 |
| PubChem | 17,520 | 11,518 | 13,750 | 4,053 | 46,841 |
| GtoPdb | 98 | 73 | 67 | 29 | 267 |
| Total | 41,972 | 26,780 | 35,204 | 11,276 | 115,232 |

**Table S4.** Trust-tier distribution for binding and functional records after curation. Gold records meet all integrity criteria; silver records meet most criteria with minor reconciliation events; exploratory records meet relaxed criteria and are reported separately rather than excluded.

| Endpoint family | Gold | Silver | Exploratory | Total |
|---|---|---|---|---|
| Binding | 14,790 | 14,070 | 920 | 29,780 |
| Functional | 4,007 | 16,592 | 598 | 21,197 |
| Total | 18,797 | 30,662 | 1,518 | 50,977 |

**Table S5.** Distribution of development-set review-queue records across uncertainty categories and review priorities. Critical and high categories are flagged for manual inspection; medium categories are tracked but allow automated downstream processing.

| Review category | Priority | Records |
|---|---|---|
| Unresolved endpoint meaning | High | 9,386 |
| Source disagreement | Critical | 4,371 |
| Source disagreement | High | 6,313 |
| Suspicious duplicate series | Medium | 1,000 |
| Pharmacology conflict | Critical | 1,003 |
| Pharmacology conflict | High | 80 |
| Suspicious duplicate activity inconsistency | High | 126 |
| Low-confidence assay semantics | Medium | 46 |

**Table S6.** Functional readout class and pharmacology class for the curated functional development records. Counts are records assigned to each label. Binding-displacement records labeled with functional pharmacology classes were routed to the binding endpoint family during endpoint matrix construction so that affinity-type measurements were not modeled as functional potency.

**Pharmacology class (functional records):**

| Pharmacology class | Records |
|---|---|
| Agonist | 13,316 |
| Antagonist | 2,704 |
| Inhibitor | 1,579 |
| Activator | 98 |
| Partial agonist | 90 |
| Allosteric modulator | 45 |
| Inverse agonist | 25 |

**Assay readout class (functional records):**

| Assay readout | Records |
|---|---|
| Binding displacement | 33,017 |
| GPCR generic / G-protein activation | 7,599 |
| cAMP | 1,501 |
| β-arrestin recruitment | 520 |
| Generic functional | 217 |
| GTPγS | 193 |
| Calcium mobilization | 146 |

**Table S7.** Internal split summary statistics for the binding and functional benchmarks under random and scaffold split regimes. Statistics are computed across model and endpoint combinations evaluated in each regime. The functional row uses the highest-resolution functional stratification, which spans the broadest endpoint set.

| Benchmark / split | Endpoint-model rows | Mean $R^2$ | Median $R^2$ | Max $R^2$ |
|---|---|---|---|---|
| Binding, random internal | 40 | 0.775 | 0.772 | 0.868 |
| Binding, scaffold internal | 40 | 0.499 | 0.615 | 0.730 |
| Functional, random internal | 320 | 0.483 | 0.590 | 0.908 |
| Functional, scaffold internal | 320 | 0.229 | 0.357 | 0.871 |

**Table S8.** Per-model external leaderboard across all 71 externally evaluated endpoints. Median and best external $R^2$ are reported per model family. The median rather than the mean is reported because a small number of low-support, high-shift functional strata produce extreme negative $R^2$ values that make the mean uninformative; these strata are visible per endpoint in Table S9. The Consensus model is the arithmetic mean of RF, XGBoost, and SVR predictions.

| Model family | Endpoints | Median external $R^2$ | Best external $R^2$ |
|---|---|---|---|
| Random Forest (RF) | 71 | 0.499 | 0.839 |
| XGBoost (XGB) | 71 | 0.490 | 0.842 |
| LightGBM (LGBM) | 71 | 0.492 | 0.846 |
| Support Vector Regression (SVR) | 71 | 0.518 | 0.808 |
| Consensus | 71 | 0.543 | 0.856 |

**Table S9.** Per-endpoint external validation across all 71 evaluated endpoints. For each endpoint the best-performing model family is reported with locked external $R^2$, RMSE, MAE, external sample size (n), the 95% bootstrap confidence interval for $R^2$ where computable, and the maximum mean discrepancy (MMD) between the training and external feature distributions. Endpoints are grouped by type (binding endpoints, then functional endpoints at increasing stratification resolution) and ordered by external $R^2$ within each group.

| Endpoint | Type | Model | $R^2$ | RMSE | MAE | n | $R^2$ 95% CI | MMD |
|---|---|---|---|---|---|---|---|---|
| pKi KOR | binding | Consensus | 0.794 | 0.669 | 0.450 | 798 | [0.75, 0.83] | 0.113 |
| pIC50 DOR | binding | XGB | 0.792 | 0.798 | 0.550 | 185 | [0.72, 0.84] | 0.186 |
| pKi DOR | binding | LGBM | 0.771 | 0.630 | 0.402 | 736 | [0.73, 0.81] | 0.111 |
| pKi MOR | binding | Consensus | 0.755 | 0.649 | 0.453 | 1227 | [0.72, 0.78] | 0.096 |
| pKi NOP | binding | XGB | 0.665 | 0.617 | 0.426 | 477 | [0.60, 0.72] | 0.159 |
| pIC50 KOR | binding | Consensus | 0.662 | 0.724 | 0.525 | 292 | [0.57, 0.74] | 0.149 |
| pIC50 MOR | binding | Consensus | 0.622 | 0.650 | 0.433 | 294 | [0.52, 0.71] | 0.141 |
| pIC50 NOP | binding | Consensus | 0.484 | 0.734 | 0.562 | 184 | [0.36, 0.58] | 0.246 |
| pIC50 func DOR | functional | RF | 0.822 | 0.599 | 0.429 | 222 | [0.77, 0.86] | 0.192 |
| pEC50 KOR | functional | XGB | 0.813 | 0.657 | 0.426 | 392 | [0.76, 0.85] | 0.138 |
| pEC50 MOR | functional | LGBM | 0.808 | 0.729 | 0.468 | 445 | [0.76, 0.85] | 0.114 |
| pEC50 DOR | functional | LGBM | 0.800 | 0.678 | 0.448 | 321 | [0.74, 0.86] | 0.132 |
| pIC50 func MOR | functional | LGBM | 0.792 | 0.573 | 0.406 | 236 | [0.72, 0.85] | 0.166 |
| pIC50 func KOR | functional | Consensus | 0.696 | 0.636 | 0.463 | 320 | [0.64, 0.75] | 0.159 |
| pEC50 NOP | functional | Consensus | 0.549 | 0.768 | 0.501 | 50 | [0.17, 0.80] | 0.310 |
| pIC50 func NOP | functional | SVR | 0.447 | 0.807 | 0.604 | 88 | [0.30, 0.56] | 0.365 |
| pAC50 MOR | functional | Consensus | 0.206 | 0.480 | 0.243 | 230 | [-0.06, 0.36] | 0.183 |
| pAC50 KOR | functional | SVR | 0.141 | 0.390 | 0.291 | 146 | [-0.04, 0.27] | 0.190 |
| pAC50 DOR | functional | Consensus | -0.014 | 0.165 | 0.116 | 143 | [-0.21, 0.10] | 0.197 |
| pIC50 func MOR antagonist | functional | Consensus | 0.856 | 0.587 | 0.449 | 110 | [0.79, 0.90] | 0.244 |
| pEC50 DOR agonist | functional | XGB | 0.811 | 0.647 | 0.428 | 308 | [0.73, 0.87] | 0.132 |
| pEC50 KOR agonist | functional | LGBM | 0.809 | 0.665 | 0.404 | 383 | [0.75, 0.85] | 0.142 |
| pEC50 MOR agonist | functional | LGBM | 0.794 | 0.739 | 0.469 | 408 | [0.75, 0.84] | 0.114 |
| pIC50 func DOR antagonist | functional | LGBM | 0.783 | 0.552 | 0.283 | 93 | [0.56, 0.91] | 0.258 |
| pEC50 DOR antagonist | functional | Consensus | 0.757 | 0.532 | 0.404 | 31 | [0.12, 0.90] | 0.226 |

| Endpoint | Type | Model | $R^2$ | RMSE | MAE | n | $R^2$ 95% CI | MMD |
|---|---|---|---|---|---|---|---|---|
| pEC50 MOR antagonist | functional | XGB | 0.733 | 0.319 | 0.195 | 70 | [-2.28, 0.91] | 0.233 |
| pIC50 func MOR agonist | functional | LGBM | 0.669 | 0.513 | 0.351 | 98 | [0.41, 0.81] | 0.283 |
| pIC50 func KOR inhibitor | functional | LGBM | 0.648 | 0.497 | 0.387 | 107 | [0.49, 0.76] | 0.300 |
| pIC50 func DOR inhibitor | functional | Consensus | 0.608 | 0.521 | 0.391 | 120 | [0.46, 0.72] | 0.273 |
| pIC50 func KOR agonist | functional | Consensus | 0.552 | 0.744 | 0.501 | 134 | [0.36, 0.67] | 0.227 |
| pEC50 NOP agonist | functional | Consensus | 0.543 | 0.803 | 0.495 | 41 | [0.13, 0.83] | 0.273 |
| pIC50 func KOR antagonist | functional | SVR | 0.518 | 0.839 | 0.636 | 146 | [0.37, 0.65] | 0.200 |
| pIC50 func MOR inhibitor | functional | LGBM | 0.426 | 0.799 | 0.525 | 33 | [-0.10, 0.82] | 0.200 |
| pAC50 MOR agonist | functional | LGBM | 0.207 | 0.482 | 0.242 | 223 | [-0.08, 0.47] | 0.183 |
| pIC50 func NOP antagonist | functional | SVR | 0.201 | 0.604 | 0.482 | 48 | [-0.37, 0.51] | 0.428 |
| pAC50 KOR agonist | functional | SVR | 0.077 | 0.355 | 0.284 | 135 | [-0.17, 0.23] | 0.197 |
| pAC50 KOR | functional | SVR | 0.052 | 0.562 | 0.491 | 34 | [-0.39, 0.17] | 0.200 |
| pAC50 MOR | functional | RF | 0.050 | 0.720 | 0.460 | 42 | [-1.06, 0.40] | 0.195 |
| pAC50 DOR agonist | functional | Consensus | -0.093 | 0.163 | 0.114 | 140 | [-0.35, 0.04] | 0.199 |
| pIC50 func MOR antagonist \| binding displacement | functional | Consensus | 0.842 | 0.497 | 0.361 | 51 | [0.71, 0.92] | 0.333 |
| pEC50 KOR agonist \| gpcr gtpgs | functional | XGB | 0.832 | 0.458 | 0.308 | 15 | [0.56, 0.94] | 0.402 |
| pEC50 KOR agonist \| binding displacement | functional | XGB | 0.764 | 0.592 | 0.376 | 180 | [0.68, 0.82] | 0.163 |
| pEC50 KOR agonist \| gpcr camp | functional | Consensus | 0.763 | 0.921 | 0.637 | 52 | [0.58, 0.86] | 0.242 |
| pEC50 DOR agonist \| gpcr generic | functional | Consensus | 0.742 | 0.764 | 0.580 | 225 | [0.67, 0.80] | 0.138 |
| pEC50 MOR agonist \| gpcr generic | functional | LGBM | 0.739 | 0.822 | 0.616 | 295 | [0.68, 0.79] | 0.120 |
| pEC50 DOR agonist \| binding displacement | functional | XGB | 0.712 | 0.592 | 0.390 | 80 | [0.56, 0.83] | 0.185 |
| pEC50 MOR agonist \| binding displacement | functional | XGB | 0.666 | 0.783 | 0.472 | 128 | [0.56, 0.77] | 0.180 |
| pEC50 KOR agonist \| gpcr generic | functional | XGB | 0.657 | 0.820 | 0.587 | 215 | [0.55, 0.74] | 0.181 |
| pIC50 func KOR antagonist \| gpcr generic | functional | SVR | 0.594 | 0.742 | 0.592 | 113 | [0.43, 0.72] | 0.232 |
| pIC50 func KOR inhibitor \| binding displacement | functional | XGB | 0.577 | 0.549 | 0.483 | 21 | [0.10, 0.74] | 0.274 |
| pIC50 func MOR antagonist \| gpcr generic | functional | XGB | 0.522 | 0.558 | 0.375 | 55 | [0.13, 0.73] | 0.400 |
| pIC50 func MOR inhibitor \| binding displacement | functional | LGBM | 0.514 | 0.741 | 0.522 | 30 | [0.15, 0.78] | 0.210 |
| pIC50 func DOR inhibitor \| binding displacement | functional | SVR | 0.492 | 0.674 | 0.547 | 38 | [0.16, 0.67] | 0.239 |
| pIC50 func KOR antagonist \| binding displacement | functional | SVR | 0.453 | 0.782 | 0.507 | 27 | [-0.06, 0.77] | 0.286 |
| pIC50 func KOR inhibitor \| gpcr generic | functional | Consensus | 0.438 | 0.477 | 0.336 | 86 | [0.16, 0.66] | 0.361 |
| pEC50 MOR agonist \| gpcr camp | functional | Consensus | 0.420 | 1.040 | 0.752 | 84 | [0.20, 0.57] | 0.218 |
| pEC50 NOP agonist \| binding displacement | functional | SVR | 0.348 | 0.868 | 0.531 | 25 | [-0.02, 0.71] | 0.271 |
| pEC50 DOR agonist \| gpcr camp | functional | SVR | 0.306 | 0.798 | 0.576 | 62 | [-0.53, 0.65] | 0.313 |
| pIC50 func NOP antagonist \| binding displacement | functional | SVR | 0.272 | 0.577 | 0.466 | 48 | [-0.23, 0.55] | 0.428 |
| pAC50 MOR agonist | functional | LGBM | 0.207 | 0.482 | 0.242 | 223 | [-0.08, 0.47] | 0.183 |
| pEC50 NOP agonist \| generic functional | functional | SVR | 0.200 | 0.996 | 0.769 | 32 | [-0.36, 0.57] | 0.310 |
| pIC50 func MOR agonist \| gpcr camp | functional | RF | 0.194 | 0.379 | 0.295 | 82 | [-0.33, 0.52] | 0.328 |
| pIC50 func DOR inhibitor \| gpcr generic | functional | SVR | 0.185 | 0.462 | 0.334 | 82 | [-0.10, 0.36] | 0.377 |
| pAC50 KOR agonist | functional | SVR | 0.077 | 0.355 | 0.284 | 135 | [-0.17, 0.23] | 0.197 |
| pAC50 KOR | functional | SVR | 0.052 | 0.562 | 0.491 | 34 | [-0.39, 0.17] | 0.200 |
| pAC50 MOR | functional | RF | 0.050 | 0.720 | 0.460 | 42 | [-1.06, 0.40] | 0.195 |
| pAC50 DOR agonist | functional | Consensus | -0.093 | 0.163 | 0.114 | 140 | [-0.35, 0.04] | 0.199 |

| Endpoint | Type | Model | $R^2$ | RMSE | MAE | n | $R^2$ 95% CI | MMD |
|---|---|---|---|---|---|---|---|---|
| pEC50 MOR antagonist \| gpcr generic | functional | Consensus | -0.192 | 0.242 | 0.165 | 68 | [-1.49, 0.01] | 0.239 |
| pIC50 func KOR agonist \| gpcr beta arrestin | functional | XGB | -0.201 | 0.599 | 0.404 | 98 | [-0.69, 0.10] | 0.296 |
| pEC50 DOR antagonist \| gpcr generic | functional | Consensus | -0.223 | 0.354 | 0.259 | 27 | [-4.93, 0.08] | 0.248 |
| pIC50 func DOR antagonist \| gpcr generic | functional | SVR | -1199.895 | 0.139 | 0.134 | 66 | [-2.658897628864375e+28, -394.98] | 0.352 |

*MMD is reported with an RBF kernel and median-heuristic bandwidth; larger values indicate greater distribution shift between training and external chemical space. Negative $R^2$ values on small, high-shift strata indicate that the model predicts worse than the endpoint mean on that external partition and are retained for transparency.*

**Table S10.** Fully pooled versus stratified functional comparison. For each receptor and activity scale, a fully pooled functional baseline (all pharmacology classes and assay readouts combined, with binding-displacement records retained in the functional pool) is compared with the best pharmacology- and readout-stratified endpoint under the same scaffold-locked external protocol. The binding-displacement fraction of the pooled set is shown; high pooled $R^2$ on functional-IC50 and AC50 endpoints coincides with high binding-displacement content, indicating the pooled model predicts affinity rather than the labeled functional activity.

| Receptor / scale | Pooled $R^2$ | Pooled n | Pool binding % | Best stratified $R^2$ | Best stratum | Strat. n |
|---|---|---|---|---|---|---|
| DOR pEC50 | 0.809 (LGBM) | 321 | 26% | 0.811 (XGB) | agonist | 308 |
| KOR pEC50 | 0.818 (XGB) | 392 | 36% | 0.832 (XGB) | agonist, GTPγS | 15 |
| MOR pEC50 | 0.829 (LGBM) | 445 | 23% | 0.794 (LGBM) | agonist | 408 |
| NOP pEC50 | 0.574 (Consensus) | 50 | 53% | 0.543 (Consensus) | agonist | 41 |
| DOR pIC50_func | 0.825 (XGB) | 330 | 69% | 0.783 (LGBM) | antagonist | 93 |
| KOR pIC50_func | 0.709 (Consensus) | 460 | 60% | 0.648 (LGBM) | inhibitor | 107 |
| MOR pIC50_func | 0.764 (Consensus) | 448 | 80% | 0.856 (Consensus) | antagonist | 110 |
| NOP pIC50_func | 0.488 (RF) | 187 | 93% | 0.272 (SVR) | antagonist, binding displacement | 48 |
| DOR pAC50 | 0.007 (Consensus) | 143 | 100% | -0.093 (Consensus) | agonist | 140 |
| KOR pAC50 | 0.270 (SVR) | 146 | 100% | 0.077 (SVR) | agonist | 135 |
| MOR pAC50 | 0.333 (LGBM) | 230 | 71% | 0.207 (LGBM) | agonist | 223 |

*For EC50 endpoints, where binding contamination is low, pooled and stratified models perform almost identically. For functional-IC50 and AC50 endpoints, the pooled set is 60 to 100 percent binding-displacement records, and the apparently strong pooled $R^2$ reflects affinity prediction rather than functional potency. The MOR functional-IC50 stratified antagonist endpoint outperforms the contaminated pool, and the all-binding AC50 pools collapse once isolated from genuine functional signal.*

**Section S4.4. SHAP export summary**

The benchmark produced 100 Random Forest SHAP summary panels, comprising 8 binding panels and 92 functional panels spanning the action- and readout-resolved functional endpoints at increasing stratification resolution. SHAP was anchored to Random Forest models so that descriptor contributions could be compared consistently across endpoints and split regimes, even where another model family produced the best external $R^2$. The main-text interpretation (Figure 5) focuses on scaffold-split panels because scaffold transfer is the stricter generalization setting. The binding panels cover receptor-level MOR, DOR, KOR, and NOP models; the functional panels cover the action- and readout-resolved tasks that anchor the main-text claims, spanning broad-support agonist endpoints and smaller antagonist and inhibitor readouts. For the Random Forest models that anchor the SHAP analysis, locked external performance on receptor pKi was 0.7744 for KOR (n = 798), 0.7558 for DOR (n = 736), 0.7445 for MOR (n = 1,227), and 0.5905 for NOP (n = 477).

### Section S5. Supporting data

The supporting data accompanying this article contains the curated datasets and the endpoint-level validation results. The principal contents are listed below.

| File | Description |
|---|---|
| curated_data_development.csv | Curated development-set measurements with standardized structure, target, endpoint, activity value and scale, assay readout, pharmacology class, physicochemical descriptors, and scaffold assignment. |
| curated_data_external.csv | Curated scaffold-locked external-set measurements with the same fields. |
| scaffold_split_assignments.csv | Scaffold-split assignments for the development and external partitions. |
| external_validation_metrics.csv | Per-endpoint external validation metrics, including $R^2$, RMSE, MAE, external sample size, bootstrap confidence intervals, and distribution-shift diagnostics (Table S9). |
| pooled_vs_stratified_comparison.csv | Fully pooled versus stratified functional comparison (Table S10). |

*The curated datasets include per-record source and reference identifiers where available, so that individual measurements can be traced to their originating reports. Endpoint columns encode receptor, activity scale, pharmacology class, and assay readout. Together with the methods described in the main text, these data are sufficient to reproduce the curated endpoints and the reported validation results.*
