## Supplementary material for "Pharmacological Stratification of Public Bioactivity Databases: A Reusable, OECD-Anchored Curation and Benchmarking Framework Demonstrated for Opioid Receptors": SI description: README.txt

Supporting Data

Pharmacological Stratification of Public Bioactivity Databases:
A Reusable, OECD-Anchored Curation and Benchmarking Framework
Demonstrated for Opioid Receptors

Contents
--------
data/
  curated_data_development.csv   Curated development-set measurements. Each row is a
                                 measurement with standardized structure (SMILES, InChIKey),
                                 target, endpoint, activity value and scale, assay readout,
                                 pharmacology class, physicochemical descriptors, and scaffold.
  curated_data_external.csv      Curated scaffold-locked external-set measurements (same fields).
  scaffold_split_assignments.csv Scaffold-split assignments for the development and external sets.

benchmark/
  external_validation_metrics.csv      Per-endpoint external validation results: R-squared,
                                       RMSE, MAE, external sample size, 95% bootstrap confidence
                                       intervals, and a distribution-shift diagnostic.
  pooled_vs_stratified_comparison.csv  Comparison of fully pooled versus stratified functional
                                       endpoints under the same external validation protocol.

Notes
-----
- Endpoint columns encode receptor, activity scale, pharmacology class, and assay readout.
- The nociceptin receptor is denoted NOP.
- Per-record source and reference identifiers are included where available so that
  individual measurements can be traced to their originating reports.
- Activity values are reported on a negative-log molar scale where applicable.
